# Contracted anterior-posterior systematic covariance of cortical thickness in early-onset schizophrenia

**DOI:** 10.1101/2024.06.03.597077

**Authors:** Yun-Shuang Fan, Yong Xu, Bin Wan, Wei Sheng, Chong Wang, Sofie Louise Valk, Huafu Chen

**Affiliations:** The Clinical Hospital of Chengdu Brain Science Institute, School of Life Science and Technology, University of Electronic Science and Technology of China, Chengdu, China; Otto Hahn Group Cognitive Neurogenetics, Max Planck Institute for Human Cognitive and Brain Sciences, Leipzig, Germany; Department of Psychiatry, First Hospital/First Clinical Medical College of Shanxi Medical University, Taiyuan, China; MOE Key Lab for Neuroinformation, High-Field Magnetic Resonance Brain Imaging Key Laboratory of Sichuan Province, University of Electronic Science and Technology of China, Chengdu, China; Institute of Neuroscience and Medicine (INM-7: Brain and Behavior), Research Centre Jülich, Jülich, Germany

**Keywords:** cortical thickness, early-onset schizophrenia, geodesic distance, structural covariance, systematic organization

## Abstract

**Background and Hypothesis:** Schizophrenia is a neurodevelopmental condition with alterations in both sensory and association cortical areas. These alterations have been reported to follow structural connectivity patterning, and to occur in a system-level fashion. Here, we investigated whether pathological alterations of schizophrenia originated from an early disruption of cortical organization by using 7−17-years-old individuals with early-onset schizophrenia (EOS).

**Study Design:** We estimated cortical thickness using T1-weighted structural MRI data from 95 patients with antipsychotic-naive first-episode EOS and 99 typically developing (TD) controls. We then computed structural covariance of cortical thickness and estimated system-level organizational axes by performing nonlinear dimensionality reduction techniques on covariance matrices for the EOS and TD groups. Finally, we tested for group differences between EOS and TD individuals in terms of both structural covariance and covariance distances along the systematic axis.

**Study Results:** The first covariance gradient differentiated motor regions from other cortical areas. Similar to the macrostructural axis in adults, the second gradient axis in young TD discriminated anterior from posterior regions and was compressed in EOS patients relative to TD controls. In addition, patients showed increased structural covariance between two ends of the systematic axis, with increased geodesic distance of covarying regions between two ends.

**Conclusion:** Our findings revealed a contracted organizational axis of cortical thickness in EOS patients, which was attributed to excessive distally coordinated changes between anterior and posterior regions of the cortex. Taken together, our study suggests a possible early disruption of system-level neurodevelopment in schizophrenia.

## Introduction

Schizophrenia is a psychiatric disorder associated with pathological changes in gray and white matter throughout the cerebral cortex ^1^. Although the behavioral manifestations of the disease usually appear in early adulthood, numerous neuroimaging studies suggest that the pathological process of the disease begins early in brain development ^2^. Early-onset schizophrenia (EOS), which is thought to be neurobiologically continuous with its adult counterpart ^3^, provides an opportunity to study disease-specific aberrations in neurodevelopmental processes. Converging evidence suggests widespread alterations in cortical thickness in EOS patients, particularly in the frontal, temporal, and parietal regions ^4, 5^. These grey matter changes have been suggested to follow the white matter organization of the cortex, consistent with models of disease propagation ^6-8^. However, it remains unclear how maturational processes of cortical thickness are coordinately disturbed by the disease.

An intuitive method for capturing coordinated changes in cortical thickness across the cortex is the “structural covariance” approach ^9^. By calculating the covariance of cross-sectional cortical thickness data, this approach measures the similarity of anatomical variations in the brain. The covariance pattern reflects the coordinated effects of specific micro- and mesoscopic factors, such as common gene expression ^10^, synaptogenesis ^11^, and laminar thickness ^12^. In addition, the structural covarying pattern of childhood-adolescence reflects synchronized developmental changes in the cortex ^9, 13^ and serves as a signature of coordinated developmental processes ^13^. For example, this structural covariance pattern has been shown to resemble intra-individual maturational coupling inferred from longitudinal data ^14^. In addition, altered structural covariance patterns are associated with a variety of mental health conditions in young patients with disorders such as depression ^15^ and anxiety symptoms ^16^. Atypical coordinated maturations of cortical thickness have also been shown in pediatric individuals at high risk for psychosis ^17^. Thus, structural covariance, which has been suggested to reflect brain maturational processes in youth, may serve as a powerful method for investigating structural coordinated changes in EOS.

Brain maturation processes have been reported to occur in a system-like manner ^18^. Systematic patterns of brain organization have been described within a framework of “gradients” ^19^, that capture an orderly spatial progression of cortical features ^20^. For example, neuronal density change systematically along spatially organized gradients ^21^. To characterize the macroscale gradient pattern of cortical organization, a nonlinear dimensionality reduction technique, also called “diffusion embedding”, has recently been proposed ^22, 23^. By embedding cortical regions into a continuous gradient map according to the similarity of their structural covariance profiles, previous work has revealed a non-random spatial organization of structural coordination across the cortex ^24, 25^. In particular, the dominant structural gradient axis tends to distinguish posterior and unimodal cortices from anterior and transmodal cortices ^24^. Furthermore, common disease effects across various psychiatric disorders have been found to follow a specific cortical thickness covariance gradient axis ^26^. Thus, system-level disruptions of structural maturation processes in EOS could be detected by measuring gradient patterns of structural covariance.

Physical distance along the cortical surface is an important determinant of how regions are connected and thus how the brain is organized ^27^. Indeed, a previous quantitative retrograde tracer analysis of macaque cortical networks suggests that physically close areas are more likely to be interconnected ^28^. However, in addition to physical distance, anatomical similarity between regions is also an important determinant ^12, 29^. For example, cortical regions with similar laminar thickness patterns have been reported to have higher structural and functional connectivity ^12^. In fact, physical distance, anatomical similarity, and brain connectivity are intrinsically linked. In particular, physically close regions tend to have similar microstructural profiles and high interregional connectivity ^30, 31^. A structural covariance network reflecting the similarity of anatomical variations in the brain has shown a bias toward short-distance connections ^13^. Physical distance along the cortical surface has been suggested to be associated with system-level transitions of both microscale cortical cytoarchitectural covariance ^12^ and macroscale cortical thickness covariance ^24^.

To investigate whether macrostructural covariance is systematically altered in EOS patients, we first computed structural covariance of cross-sectional cortical thickness data in 7−17-years-old individuals with EOS and in typically developing (TD) controls ^13^. We then decomposed the similarity matrix of covariance into a low-dimensional embedding using the diffusion embedding approach ^22^. Based on previous work ^24^, we expected that a structural gradient would consistently describe an anterior-posterior transition, and could be distorted by EOS. To further unravel the potential biological mechanisms behind system-level changes of cortical thickness in patients, we examined disease effects on structural covariance patterns ordered by the covariance gradient axis in TD controls. Finally, we evaluated the relationships between geodesic distances and gradient axes, and estimated covariance distances by calculating geodesic distance between covarying regions ^32^. In the case of a disruption of cortical organization in EOS, we expect that covariance distance to be affected by the disease.

## Material and methods

### Participants

Ninety-nine drug-naive first-episode EOS patients and 100 TD controls were recruited from the First Hospital of Shanxi Medical University, Taiyuan, China. The diagnosis of schizophrenia was made according to the Structured Clinical Interview for Diagnostic and Statistical Manual of Mental Disorders, Fourth Edition, and confirmed by at least one senior psychiatrist (Y.X.). The psychiatric symptomatology of 71 patients was assessed using the Positive and Negative Syndrome Scale (PANSS). The absence of the remaining patients was due to their relatively low compliance. Exclusion criteria for all subjects included i) age over 18 years; ii) history of neurological MRI abnormalities; iii) substance abuse; and iv) any electronic or metal implants. Patients were also excluded if they had had the disease for > 1 year. TD controls were also excluded if they had a personal or family history of psychiatric disorders. This study was approved by the Ethics Committee of the First Hospital of Shanxi Medical University. Informed consent was obtained from all participants and their parents or legal guardians.

### Image data acquisition

T1-weighted anatomical data were collected using a 3 Tesla Siemens MAGNETOM Verio scanner at the First Hospital of Shanxi Medical University. Image data were acquired via a three-dimensional fast spoiled gradient-echo sequence. Scanning parameters included the following: repetition time = 2,300 ms, echo time = 2.95 ms, flip angle = 9°, matrix = 256 × 240, slice thickness = 1.2 mm (no gap), and voxel size = 0.9375 × 0.9375 × 1.2 mm^3^, with 160 axial slices.

### Cortical thickness extraction

Anatomical images were first preprocessed using the FreeSurfer package (version 7.1.0, http://surfer.nmr.mgh.harvard.edu/) ^33^, including cortical segmentation and surface reconstruction. Four patients were excluded due to incomplete scanning and one control due to poor quality of cortical parcellation, resulting in a final sample including 95 EOS patients and 99 demographically-matched TD controls (**Table 1**). Vertex-wise cortical thickness values were then estimated using the distance between the white and pial surfaces. Subsequently, surface vertices were down-sampled to 400 cortical parcels via “Schaefer” local-global atlas ^34^. Parcel-wise cortical thickness was estimated by averaging vertex-wise thickness values within each parcel. Consistent with previous findings ^35^, patients with EOS (mean ± SD = 2.58 ± 0.45 mm) showed reduced global cortical thickness relative to TD controls (mean ± SD = 2.62 ± 0.46 mm; *t* = 2.41, *p* = 0.02).

**Table 1.**
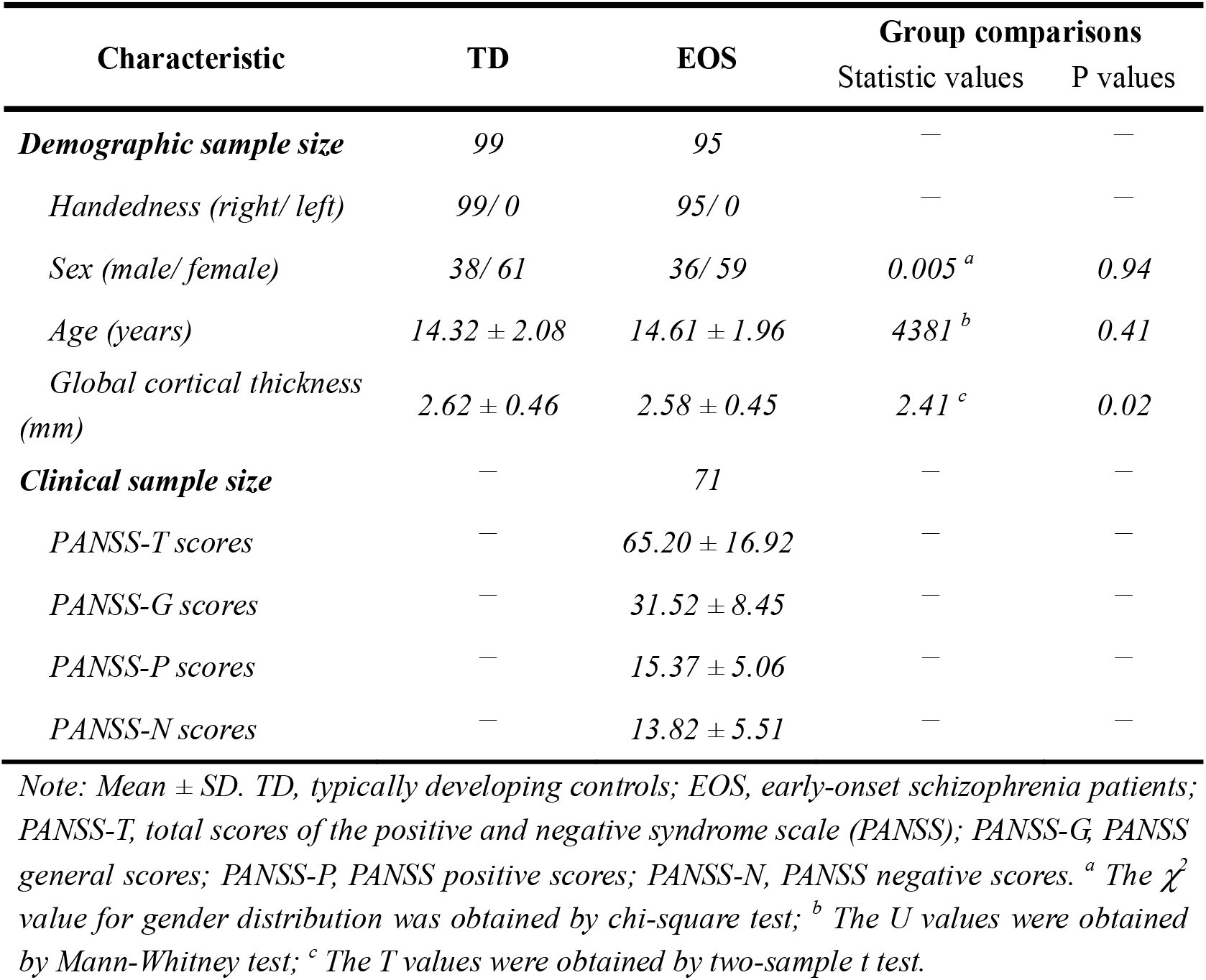
Demographic and clinical characteristics.

### Structural covariance gradient calculation

To investigate system-level structural covariance patterns, we computed the structural covariance matrix separately for the EOS and TD groups. Specifically, we computed partial Pearson’s correlations between each pair of cross-sectional cortical thickness data with sex, age, and global cortical thickness as covariates. We then performed Fisher’s z-transformation on the correlation matrix. We then estimated systematic covariance gradients using the BrainSpace toolbox (https://github.com/MICA-MNI/BrainSpace) ^36^. Briefly, the *z*-transformed covariance matrix was column-wise thresholded at 90% and transformed into an affinity matrix by using a normalized angle similarity kernel. Next, its dimensionality was nonlinearly reduced by using a diffusion embedding method (α = 0.5, a parameter which controls the impact of sampling density) ^22, 37^. The gradient maps for the EOS group were aligned with a normative gradient mask generated by the TD group using Procrustes rotations. Along the continuous gradient axis, close gradient scores of two regions reflected similar structural covarying profiles. We also estimated covariance gradient maps by using only sex and age as covariates, and found similar gradient patterns as the original gradient maps (**Figure S1**).

To obtain the gradient scores with statistical parameters, we transformed gradient scores to *z-*scores. We then calculated group-level differences in *z* scores ^38^ and corrected them using the false discovery rate (FDR) method (*q*_*FDR*_ < 0.05) to assess statistical significance. The nonparametric Kolmogorov-Smirnov test was used to compare the distributions of gradient scores between the TD and EOS groups ^39^. In addition, to characterize the functional involvement of covariance gradients, we further grouped 400 cortical parcels into seven functional networks, including the visual network (VIS), sensorimotor network (SMN), dorsal attention network (DAN), ventral attention network (VAN), limbic network (LMB), frontoparietal network (FPN), and default mode network (DMN) ^40^. Network-level group differences between patients and controls were examined by performing paired *t*-tests on gradient scores of all parcels belonging to a given network (FDR corrected, *q*_*FDR*_ < 0.05).

### Disease effects on structural covariance

To further determine systematic covariance changes in EOS patients, we examined disease effects on structural covariance patterns along the system-level gradient axis. First, we divided cortical parcels into 10 equal-sized bins according to their ranked gradient scores in TD controls. We then averaged the structural covariance matrices within these bins, resulting in 10 × 10 covariance matrices. Next, we examined diagnostic effects by using a classical linear interaction model with diagnosis and cortical thickness as two factors, as implemented in BrainStat (https://github.com/MICA-MNI/BrainStat) ^41^. Finally, we examined symptom effects in the EOS group by using PANSS positive (or negative) scores and cortical thickness as the two factors. In these linear models, sex, age, and global cortical thickness were regressed out, and FDR corrections (*q*_*FDR*_ < 0.05) were used to control for the effect of false positives.

### Relationships with geodesic distance

To evaluate the relationship between systematic structural covariance and physical distance, we first computed the geodesic distance matrix across the cortex. The geodesic distance between two parcels refers to the length of their shortest path on the mesh-based representation of the cortex. Specifically, we computed the geodesic distance between each vertex in fsaverage5 space, and then took the average distance between both parcels to obtain parcel-wise distances by using the Micapipe toolbox (https://micapipe.readthedocs.io/) ^42^. The geodesic distance was first calculated within each hemisphere, and then averaged across two hemispheres to represent the interhemispheric geodesic distance. We then calculated Pearson’s correlations between the gradient maps in TD and the node-wise degree map of geodesic distance. Statistical significance (p_spin_ < 0.05) was estimated by using the spin test implemented in the ENIGMA toolbox (https://enigma-toolbox.readthedocs.io/en/latest/) ^43, 44^. The spin test simulates 10,000 surrogate surface maps with spatial autocorrelation and generates a null distribution of correlation values. We then computed Pearson’s correlation between the structural covariance matrix in TD and the edge-wise geodesic distance matrix, and assessed the significance by using permutation test (*p*_*perm*_ < 0.05, 10,000 times).

### Disease effects on covariance distances

To further elucidate system-level covariance changes in EOS, we estimated disease effects on covariance distances. Covariance distance refers to the averaged geodesic distance from a seed region to its structurally covarying regions ^32^. Specifically, a column-wise 90% -threshold structural covariance matrix was used as a mask to average the geodesic distance profiles of each parcel to generate the covariance distance map. A high covariance distance score of a seed region reflected a distant projection pattern, while a low score indicated a local projection. Group-level differences between covariance distances of TD and EOS were measured by comparing *z*-score maps of the covariance distance between two groups. In addition, we tested for disease effects by performing paired *t*-tests on covariance distance scores of all parcels belonging to a given gradient bin (*q*_*FDR*_ < 0.05).

## Results

To reveal system-level structural abnormalities of EOS patients, we calculated structural covariance gradients of cortical thickness for the TD and EOS groups (**Figure 1**). Specifically, we decomposed the 90% thresholded covariance matrix into 10 gradient components and aligned gradient maps of EOS with normative gradient maps of TD controls. Along the gradient axis, the position of a region was determined by the similarity of its structural covariance profile to others, thus indicating opposite poles of the axis with maximally divergent covariance patterns. The first gradient component (G1) explained 20% of the variance in the TD group and 26% of the variance in the EOS group to distinguish motor regions from other cortical areas. However, only the second gradient (G2) axis (explained eigenvariance: EOS group = 18%; TD group = 15%) was significantly related to the main gradient axis from the Human Connectome Project (HCP) data (*r* = 0.69, *p*_*spin*_ < 0.0001) ^24^, whereas the G1 map showed no correlation with either the HCP G1 map (*r* = 0.22, *p*_*spin*_ = 0.18) or the HCP G2 map (*r* = 0.20, *p*_*spin*_ = 0.28) (**Figure S2**).

**Figure 1.**
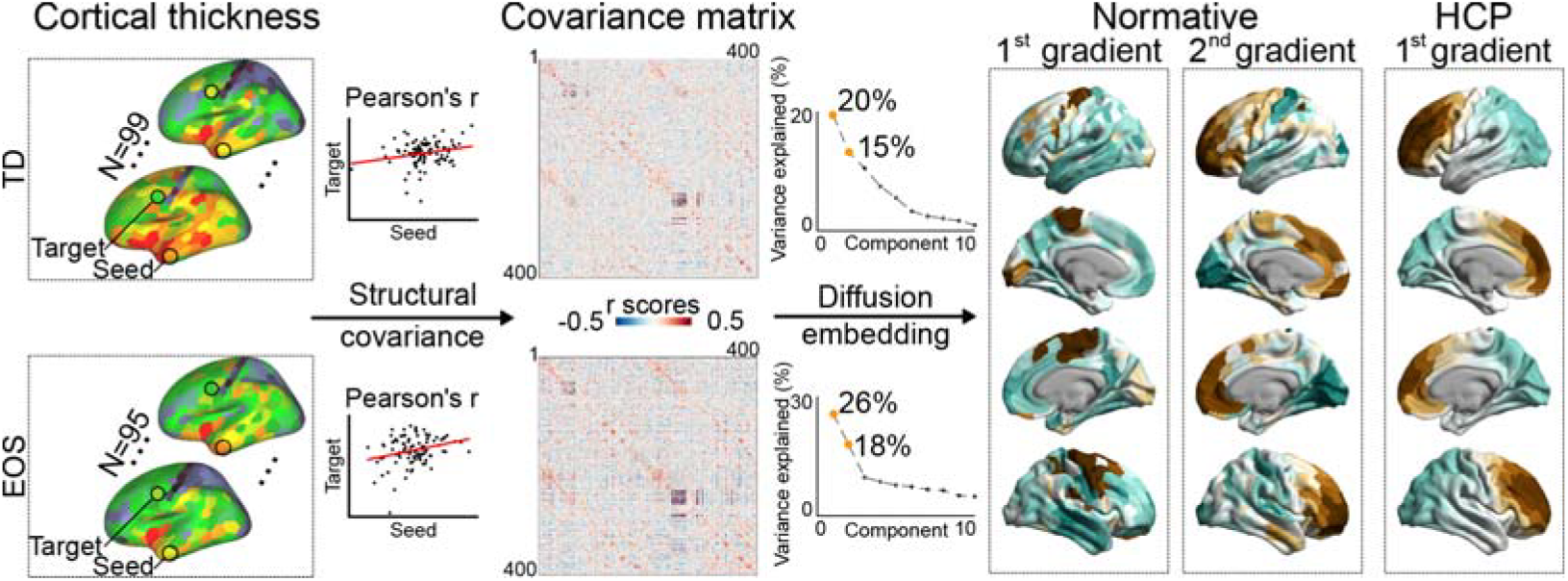
Method flowchart of the structural covariance gradient. Vertex-wise cortical thickness was first extracted and down-sampled to 400 parcels from the Schaefer atlas ^34^. Structural covarying patterns were then estimated by using structural covariance of cortical thickness in typically developing (TD) controls, or early-onset schizophrenia (EOS) patients. After column-wise thresholding at 90%, the covariance matrix was then decomposed into 10 low dimensional components by using the diffusion embedding method. The second component in TD, i.e., the second normative gradient, was similar with the first gradient axis derived from the human connectome project (HCP) data ^24^, and thus selected-out and used in further analyses.

### Systematic covariance gradient maps

The G2 axis, which is similar to the HCP principal gradient map ^24^ described a spatial arrangement from anterior to posterior regions in the cerebral cortex (**Figure 2A**). Patients with EOS showed a compressed gradient axis compared to TD controls (Kolmogorov-Smirnov test; *D*_*400*_ = 0.13, *p* = 0.002). Although no parcels survived FDR correction for z score differences of gradient scores, frontal regions showed subtly decreased gradient scores in EOS patients relative to TD controls (*p*_*uncorrected*_ < 0.05). After pooling 400 cortical parcels into seven functional networks, we observed that the posterior end was located in the VIS and SMN, while the anterior end was anchored in transmodal networks including the DMN and FPN (**Figure 2B**). No significant group difference in network-level gradient scores was found between the EOS and TD groups using paired t-tests. In addition, group comparisons of the G1 map suggested an extended axis in EOS patients compared to TD controls (*D*_*400*_ = 0.19, *p* < 0.0001) (See **Figure S3** for details).

**Figure 2.**
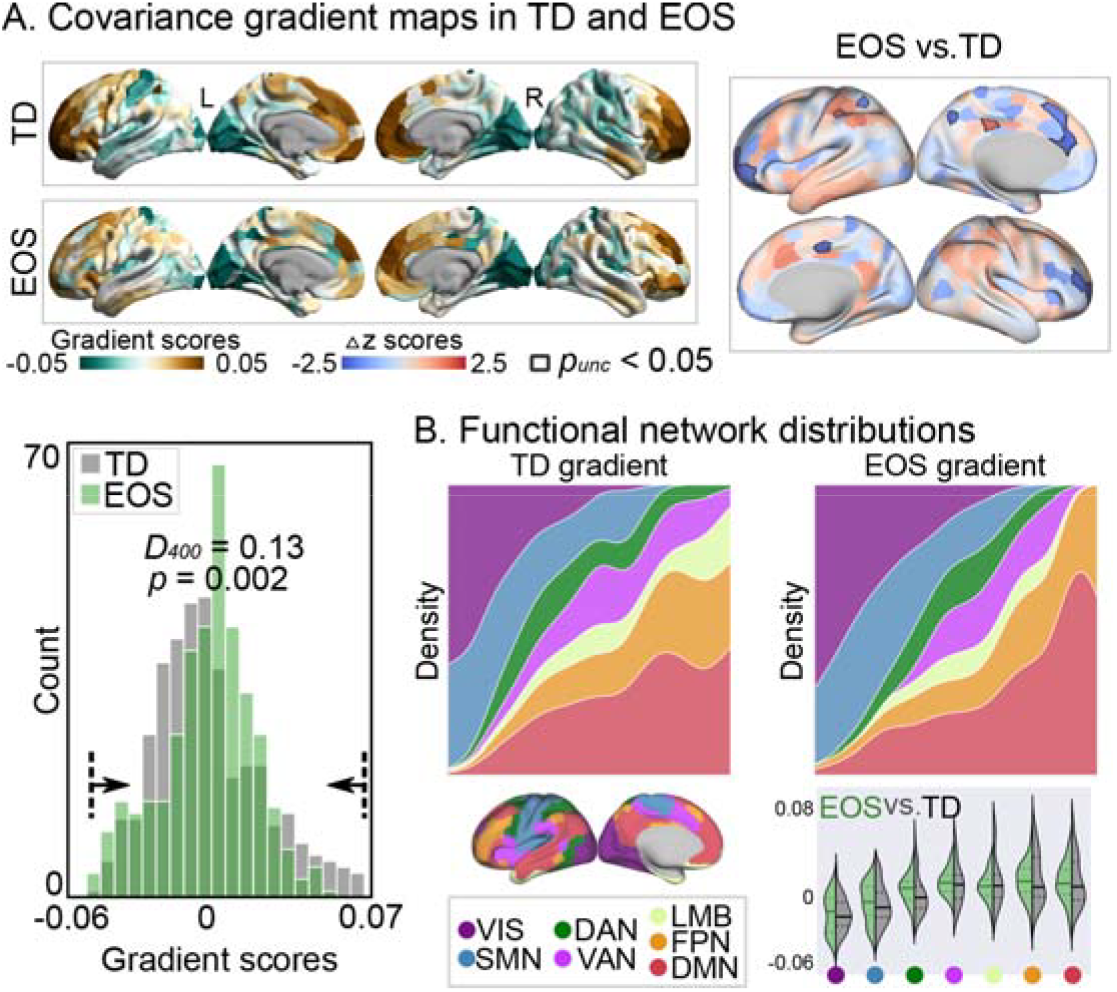
Structural covariance gradient patterns. **(A)** Covariance gradient maps in TD and EOS. The top right brain map shows the group differences of *z* scores transformed from gradient scores. Parcels showing subtle group differences (*p*_*uncorrected*_ < 0.05) are outlined in black. In the bottom density map, green boxes represent gradient scores of EOS patients and grey boxes represent TD controls, which suggested compressed gradient axis in EOS than TD (Kolmogorov-Smirnov test; *D*_*400*_ = 0.13, *p* = 0.002). **(B)** Network distributions. For each group, the continuous density maps of gradient scores were separately plotted for each network. Group differences between EOS and TD were tested by using paired *t*-tests. VIS, visual network; SMN, sensorimotor network; DAN, dorsal attention network; VAN, ventral attention network; LMB, limbic network; FPN, frontoparietal network; DMN, default mode network.

### Group differences on structural covariance

To further investigate the compressed anterior-posterior axis in EOS patients, we reorganized and averaged the structural covariance matrices according to the 10-binned normative gradient mask (**Figure 3A**). As expected, the further apart two bins were along the gradient axis, the lower the covariance value. We then tested for diagnostic effects on the reshaped structural covariance patterns. We found that patients had increased covariance values between the 1^st^ and 9^th^ bins (*t* = 3.39, *q*_*FDR*_ = 0.004), i.e., less negative covariance values in EOS compared to TD (**Figure 3B**). Patients also showed decreased covariance values between the 7^th^ and 10^th^ bins (*t* = −2.72, *q*_*FDR*_ = 0.03), i.e., less positive covariance values in EOS patients relative to TD controls.

**Figure 3.**
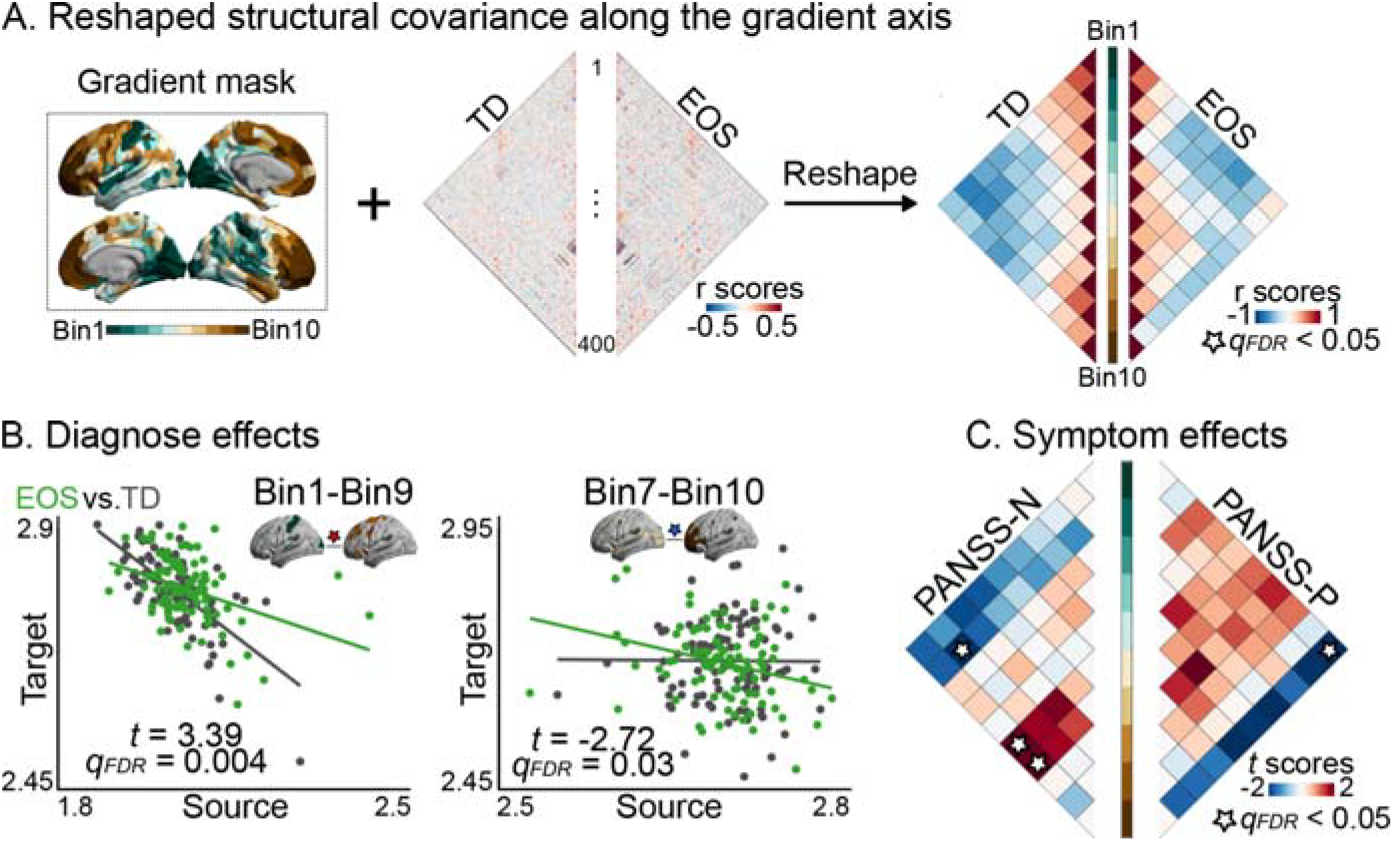
Disease effects on structural covariance patterns. **(A)** Reshaped covariance matrices along the gradient axis. Covariance matrices were reshaped into 10 × 10 matrices according to a 10-binned mask generated from the anterior-posterior gradient of TD controls. **(B)** Diagnose effects on reshaped covariance. The diagnose effect was then examined by using a classic interaction linear model with group and cortical thickness as factors [False discovery rate (FDR) corrections, *q*_*FDR*_ < 0.05]. Green dots and lines represent EOS patients and grey dots and lines represent TD controls. **(C)** Symptom effects on reshaped covariance. We used N scores of the Positive and Negative Syndrome Scale (PANSS) to quantify the severity of negative symptoms, and PANSS-P scores for positive symptoms. In the EOS group, symptom effects were examined by the interaction linear model with PANSS-P/N scores and cortical thickness as factors.

In the EOS group, we found significantly negative effects of PANSS negative scores on covariance between the 2^th^ and 9^th^ bins (*t* = −2.55, *q*_*FDR*_ = 0.03), and positive effects on covariance between the 10^th^ and 6^th^ bins (*t* = 2.29, *q*_*FDR*_ = 0.04), and the 10^th^ and 7^th^ bin (*t* = −2.5, *q*_*FDR*_ = 0.04) (**Figure 3C**). Similarly, PANSS positive scores had significantly negative effects on covariance between the 1^st^ and 10^th^ bins (*t* = −2.83, *q*_*FDR*_ = 0.01).

### Group differences on covariance distances

As shown in **Figure 4A**, we computed node-wise geodesic distance degrees by averaging geodesic distances from one parcel to all other parcels, and found that it was relatively increased in the frontal, inferior temporal, and occipital regions compared to other regions. The anterior-posterior gradient map (G2) was significantly related to the nodal degree map of geodesic distance (*r* = 0.36, *p*_*spin*_ = 0.04). However, the G1 map showed no correlation with geodesic distance (**Figure S4**). As expected, structural covariance was negatively correlated with geodesic distance (*r* = −0.17, *p*_*perm*_ = 0.001), supporting previous findings of a short-distance connection bias in structural covariance networks ^13^.

**Figure 4.**
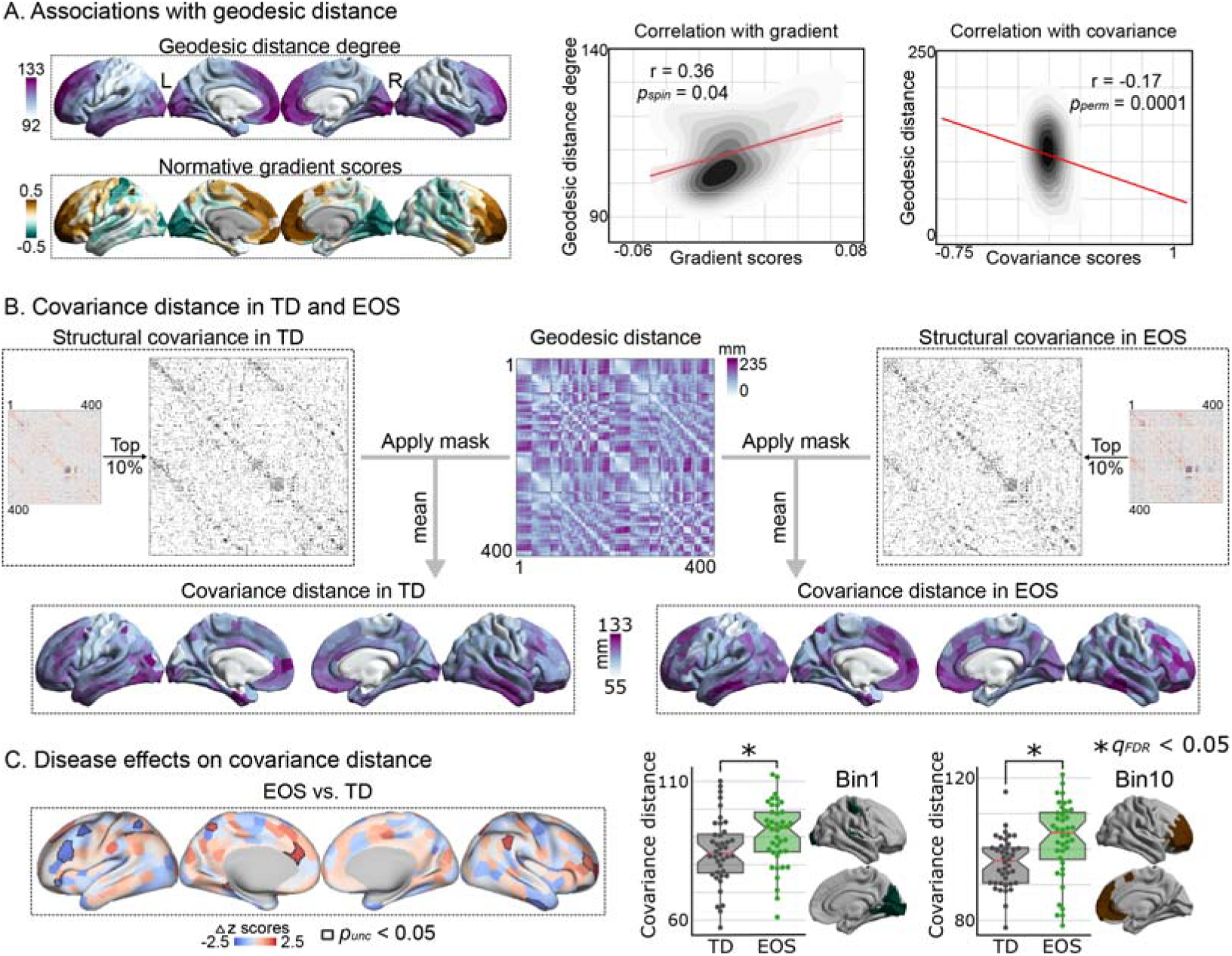
Covariance distance in TD and EOS. **(A)** Correlations between geodesic distances and covariance gradients and covariance matrices. The node-wise geodesic distance degree was calculated by averaging the geodesic distance for each node. The spin test was used to control for spatial autocorrelations (*p*_*spin*_ < 0.05, 10,000 times) ^44^. The edge-wise geodesic distance was then correlated with the structural covariance matrix, and the statistical significance was assessed using permutation tests (*p*_*perm*_ < 0.05, 10,000 times). **(B)** Covariance distance calculation. For each group, we column-wise thresholded the structural covariance matrix at 90%, and then binarized it as a mask. Covariance distance was then calculated using the averaging geodesic distance profiles of a seed region multiplied by the mask. **(C)** Effects of disease on covariance distances. In the left z-score brain map, regions with subtle group differences in *z* scores (*p*_*uncorrected*_ < 0.05) are outlined in black. In the right box-and-whisker plots, the red horizontal lines in the box indicate the mean of each group, and the lower and upper horizontal lines indicate the lower and upper quartiles, respectively. Group differences between the EOS and TD groups were tested using paired *t*-tests (* represents *q*_*FDR*_ < 0.05). Green boxes represent covariance distances of EOS patients and grey boxes represent TD controls.

To investigate the relationship between physical distances and gradient perturbations in EOS, we computed covariance distances by averaging the geodesic distance from a seed region to its covarying regions (**Figure 4B**) ^32^. A high covariance distance score of a region indicates a pattern dominated by remote connectivity, and a low score indicates local connectivity. Patients with EOS (mean ± SD = 90.15 ± 13.08 mm) had numerically increased covariance distances compared to TD controls (mean ± SD = 88.77 ± 11.93 mm). In our exploratory analyses comparing *z*-scored covariance distances of EOS and TD (**Figure 4C**), we found that patients had subtly decreased covariance distances in the left frontal regions, while there were increased distances in the right frontal regions (*p*_*uncorrected*_ < 0.05). After resampling the covariance distance maps along the gradient axis, we found that patients showed increased covariance distances in the 1^st^ bin (*t* = 3.71, *q*_*FDR*_ = 0.007) and the 10^th^ bin (*t* = 3.27, *q*_*FDR*_ = 0.01) compared to TD controls.

## Discussion

In the current study, we investigated the system-level organization of coordinated structural changes in EOS patients by applying a dimensional reduction approach to the structural covariance of cross-sectional cortical thickness data. G1 in motor regions differed from other cortical areas. Notably, similar to the principal gradient based on cortical thickness from the young adult HCP sample, the G2 of the structural covariance pattern described an anterior-posterior organizational axis, capturing a unimodal-transmodal transition. Overall, patients with EOS showed a contracted anterior-posterior gradient pattern compared to TD controls. In addition, patients showed increased structural covariance between the anterior and posterior ends of the gradient and increased covariance distances of both poles compared to TD controls. Taken together, these findings revealed a disrupted systematic organization of structural covariance patterns in EOS patients, which was supported by excessive distant connection profiles between two ends of the axis.

Consistent with previous findings ^24^, our study revealed an anterior-posterior gradient axis of structural covariance in this young age group. However, it was the second, and not the first, covariance gradient of this age group that was aligned with the main covariance gradient in adults ^24^. The shift in macroscale cortical organization between pediatric and adult populations has been reported in functional connectivity gradients ^45^. This previous work concluded that these gradient order changes represent a maturation of cortical organization that may be critical for refinement of cognitive and behavioral abilitie during development. As previously suggested ^24^, the anterior-posterior axis appears to map the temporal sequence of neurogenesis. Specifically, the posterior and anterior portions of the cortex are distinguished by their neuronal counts, i.e., a greater number of neurons with a shorter cell cycle at the posterior end, and a smaller number with a longer cell cycle at the anterior end ^46^. Furthermore, this gradient axis has been suggested to be related to the functional continuum from basic perception to abstract cognition ^24^. Our findings of functional network distributions showed a similar unimodal-transmodal transition, supporting previous gradient findings in adults.

The system-level covariance gradient axis was compressed in EOS patients, consistent with previous findings of functional gradient compression in schizophrenia ^47^. A previous study found compression of the sensorimotor-to-transmodal functional connectivity gradient in patients with chronic adult-onset schizophrenia, and suggested that this was a system-level substrate underlying sensory and cognitive deficits of patients ^48^. However, the compression of the functional gradient was mainly manifested in sensorimotor systems, whereas the macrostructural compression was located in frontal regions. Indeed, frontal regions have been suggested as the putative pathologic origin for brain tissue volume loss ^6^ or cortical thinning ^8^ in schizophrenia. Assuming that brain structure supports intrinsic function ^49^, we reasoned that impaired structural organization may play a role in functional system dysfunctions in schizophrenia. Future work using longitudinal data is needed to further elucidate their possible causal relationships.

In addition to the cerebral cortex, subcortical nuclei are also important pathological components in schizophrenia ^50^. We additionally computed a covariance gradient by combining cortical areas and subcortical regions, including the accumbens, amygdala, caudate, hippocampus, pallidum, putamen, and thalamus (**Figure S5**). We found that the cortical gradient maps were similar to the original gradient maps, showing a compressed anterior-posterior gradient axis in patients. EOS patients showed decreased covariance between the left putamen (one of the basal ganglia nuclei and part of the striatum) and the middle part of the anterior-posterior axis. In a previous patient case, a left putamen infarct was reported cause psychotic symptoms ^51^. However, it is the thalamocortical connectivity that has been shown to strongly contribute to the formation of key characteristics of the mature brain during youth ^52^. Our previous work using the same dataset found increased segregation of macroscale thalamocortical functional organization in EOS ^53^. The current coarse resolution of subcortical nuclei may account for the inconsistent findings of the thalamus, and future work examining the cerebellum and finer subcortical regions is highly recommended.

Patients showed increased structural covariance between posterior and anterior regions, which is partially consistent with previous functional findings of hyperconnectivity between unimodal and fronto-parietal regions in schizophrenia ^47^. The previous study suggested that reduced functional separation between primary sensory and fronto-parietal cognitive systems may contribute to the phenomenon of functional hierarchical compression. During brain development, primary sensory cortices are relatively uncoupled from the rest of the cortex ^9^, whereas frontotemporal cortices have stronger and more extensive coupling patterns, responsible for the involvement of integrative cognitive processes ^54^. Therefore, patients’ excessive structural coupling between posterior and anterior regions may provide a mechanistic explanation for compressed structural organization in EOS. In addition, we found that the clinical severity of both positive and negative symptoms increased with the weakening of structural coupling between posterior and anterior regions, suggesting a compensatory neural mechanism in EOS patients. However, the current study design was based on a cross-sectional dataset, that does not reflect co-maturation processes between different cortical regions in young individuals. Although population-based structural covariance of cortical thickness could be explained by subject-based maturational coupling patterns derived from longitudinal data ^14^, further longitudinal studies are needed to validate the system-level perturbations of structurally coordinated changes in EOS patients.

Geodesic distance was correlated with the anterior-posterior covariance gradient, supporting the hypothesis that physical distance is an important determinant of cortical organization. Indeed, we found that regions with greater geodesic distance had lower structural covariance. However, we again found distance-related differences between individuals with EOS and TD. By calculating the covariance distance, we tested the relationship between physical distance and systematic structural organizational changes in EOS. In general, sensory areas had more clustered local connections, whereas transmodal systems had distributed remote connections ^55-57^. The increased connectivity distance in association areas relative to sensory areas could be driven by multiple factors, reflecting a systematic balance between short- and long-distance connections. The “tethering hypothesis” relates this distribution of connectivity distance to evolutionary expansion ^58^. Specifically, this hypothesis views sensory regions as anchors and transmodal association cortex as the evolutionally expanding cortical areas tethering these anchors, potentially explaining the increase in long-range connectivity in association regions. We found that patients with EOS showed increased covariance distances of both systems, which is consistent with previous findings of distance-dependent miswiring patterns ^59^. According to a concept of network attributes ^60^, local connections are associated with functional system segregation, and long-range connections with integration. Therefore, increased covariance distances in EOS patients might reflect disturbed network topology, potentially interpreting their contracted macroscale structural organization.

## Conclusions

The current study described a contracted anterior-posterior organization of structural covariance patterns in EOS patients, which may be related to increased distant coordinated changes between posterior regions (including sensorimotor networks) and anterior regions (including transmodal networks such as the DMN and FPN). More broadly, this study provided a systems-level perspective on early structural perturbations in schizophrenia, further supporting the neurodevelopmental hypothesis.

## Data availability

Structural covariance patterns and systematic gradient maps in TD and EOS and other data supporting the findings of this study are available at https://github.com/Yun-Shuang/Structural-covariance-gradient-SZ.

## Code availability

Custom code was made publicly available under https://github.com/Yun-Shuang/Structural-covariance-gradient-SZ. Gradients calculation is based on BrainSpace (https://brainspace.readthedocs.io/en/latest/). Statistically analyses were performed using BrainStat (https://github.com/MICA-MNI/BrainStat) and ENIGMA (https://enigma-toolbox.readthedocs.io/en/latest/). Visualizations were based on workbench (https://www.humanconnectome.org/software/connectome-workbench) combined with ColorBrewer (https://github.com/scottclowe/cbrewer2).

## Acknowledgements

We are grateful to all the participants and their guardians in this study. We thank International Science Editing (http://www.internationalscienceediting.com) for editing this manuscript. This work was supported by STI 2030—Major Projects 2022ZD0208900, the National Natural Science Foundation of China (62333003, 62036003, 82121003, 62373079, 62073058), Medical-Engineering Cooperation Funds from University of Electronic Science and Technology of China (ZYGX2021YGLH201). Y-S.F. was also funded by the China Postdoctoral Science Foundation (2023M740524) and Sichuan Province Innovative Talent Funding Project for Postdoctoral Fellows. S.L.V. was also funded in part by Helmholtz Association’s Initiative and Networking Fund under the Helmholtz International Lab grant agreement InterLabs-0015, and the Canada First Research Excellence Fund (CFREF Competition 2, 2015-2016) awarded to the Healthy Brains, Healthy Lives initiative at McGill University, through the Helmholtz International BigBrain Analytics and Learning Laboratory (HIBALL). B.W. is supported by the International Max Planck Research School on Neuroscience of Communication: Function, Structure, and Plasticity (IMPRS NeuroCom).

