## Supplementary Figures for "Contracted anterior-posterior systematic covariance of cortical thickness in early-onset schizophrenia"

Running title: Early structural gradient disruptions in schizophrenia

Yun-Shuang Fan ^1,2^, Yong Xu ^3^, Bin Wan ^2^, Wei Sheng ^1^, Chong Wang ^1^, Sofie Louise Valk ^2,5*#^, Huafu Chen ^1,4*#^

*1.The Clinical Hospital of Chengdu Brain Science Institute, School of Life Science and Technology, University of Electronic Science and Technology of China, Chengdu, China; 2. Otto Hahn Group Cognitive Neurogenetics,* *Max Planck Institute for Human Cognitive and Brain Sciences, Leipzig, Germany; 3. Department of Psychiatry, First Hospital/First Clinical Medical College of Shanxi Medical University, Taiyuan, China; 4. MOE Key Lab for Neuroinformation, High-Field Magnetic Resonance Brain Imaging Key Laboratory of Sichuan Province,* *University of Electronic Science and Technology of China, Chengdu, China.* *5. Institute of Neuroscience and Medicine (INM-7: Brain and Behavior), Research Centre Jülich, Jülich, Germany.*

* Both last co-authors contributed equally.

\# Corresponding authors:

Huafu Chen, & Sofie Louise Valk,.

**Supplementary Figures**

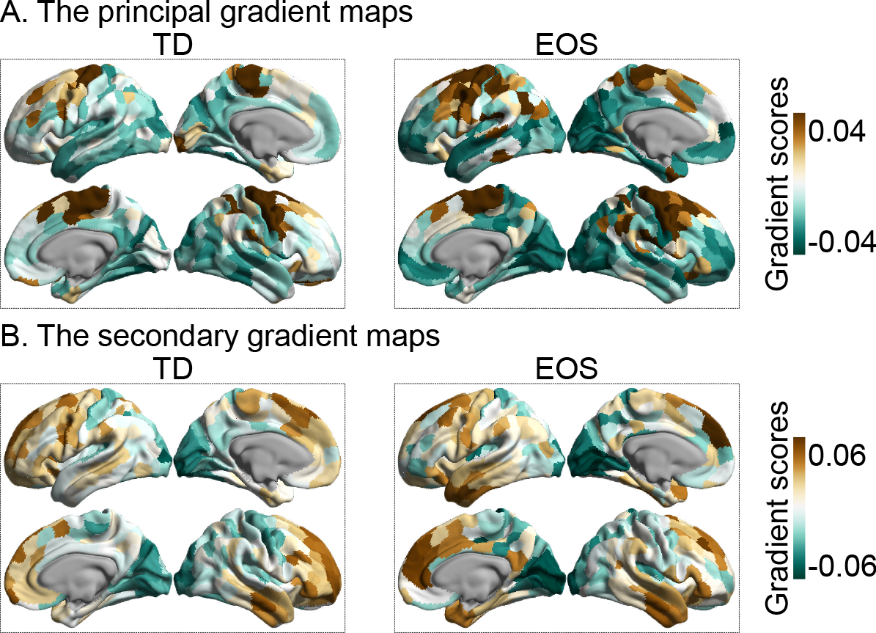

**Figure S1. Replicated covariance gradient maps.** Early-onset schizophrenia, EOS; typically developing, TD.

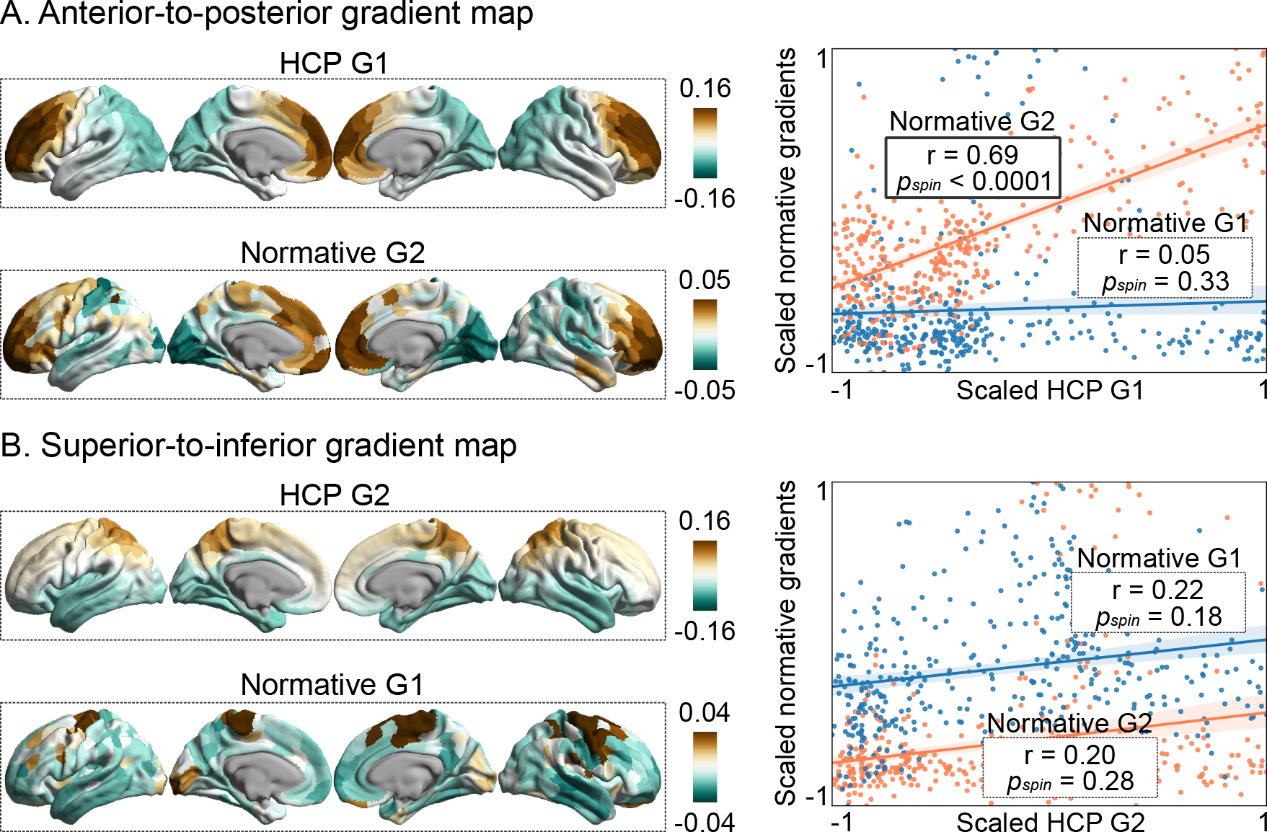

**Figure S2. Relationships with gradient maps from human connectome project (HCP) data.** **(A)** The first gradient (G1) map of HCP data showed an anterior-posterior transition, which was significantly correlated with the second gradient (G2) map of TD (*r* = 0.69, *p_spin_* < 0.0001). **(B)** The HCP G2 map described a spatial arrangement from superior cortices to inferior cortices, while it showed no significant correlation with our gradient maps.

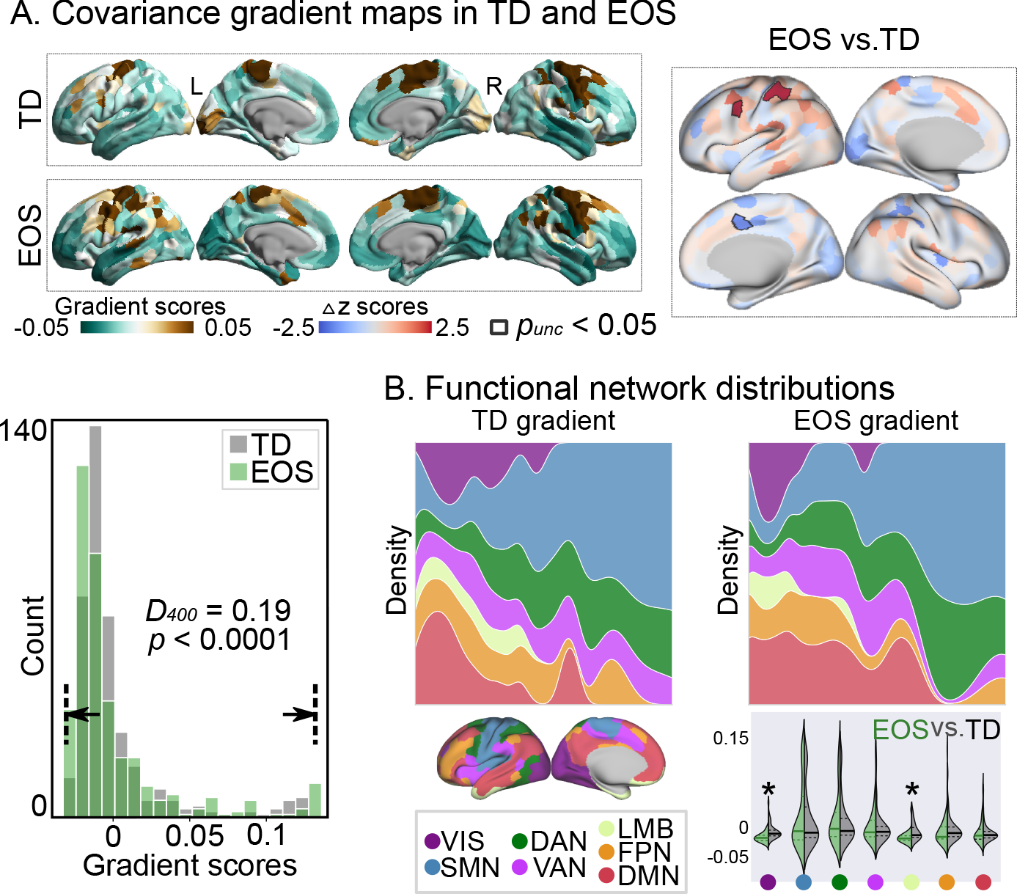

**Figure S3. The first gradient of the structural covariance.** (A) The first gradient distinguished sensorimotor regions from other regions (explained variance: EOS group = 26%; TD group = 20%). As shown in the right *z*-score map, gradient scores in sensorimotor regions were subtly increased in EOS than in TD (*p_uncorrected_* < 0.05). Patients with EOS showed an extended gradient axis compared to TD controls (Kolmogorov-Smirnov test; *D_400_* = 0.19, *p* <0.0001). (B) From a visual perspective, the first gradient differed between the visual network (VIS) and the sensorimotor network (SMN). Compared to TD controls, patients showed decreased network level gradient scores in VIS (*t* = −4.2, *q_FDR_* = 0.0006) and limbic network (LMB; *t* = −3.28, *q_FDR_* = 0.01). Dorsal attention network, DAN; ventral attention network, VAN; frontoparietal network, FPN; default mode network, DMN.

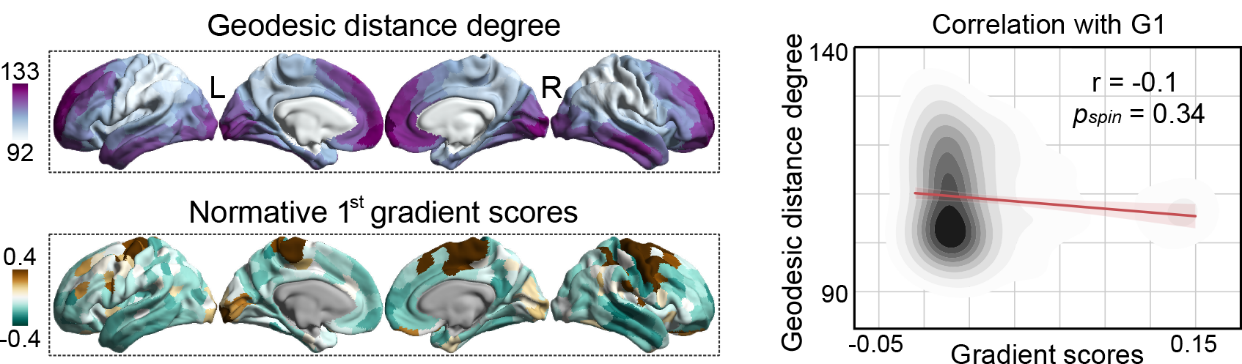

**Figure S4. Association between geodesic distance degree and the first covariance gradient.** No significant correlation was found between normative G1 map and node-wise geodesic distance degree map (*r* = −0.1, *p_spin_* = 0.34).

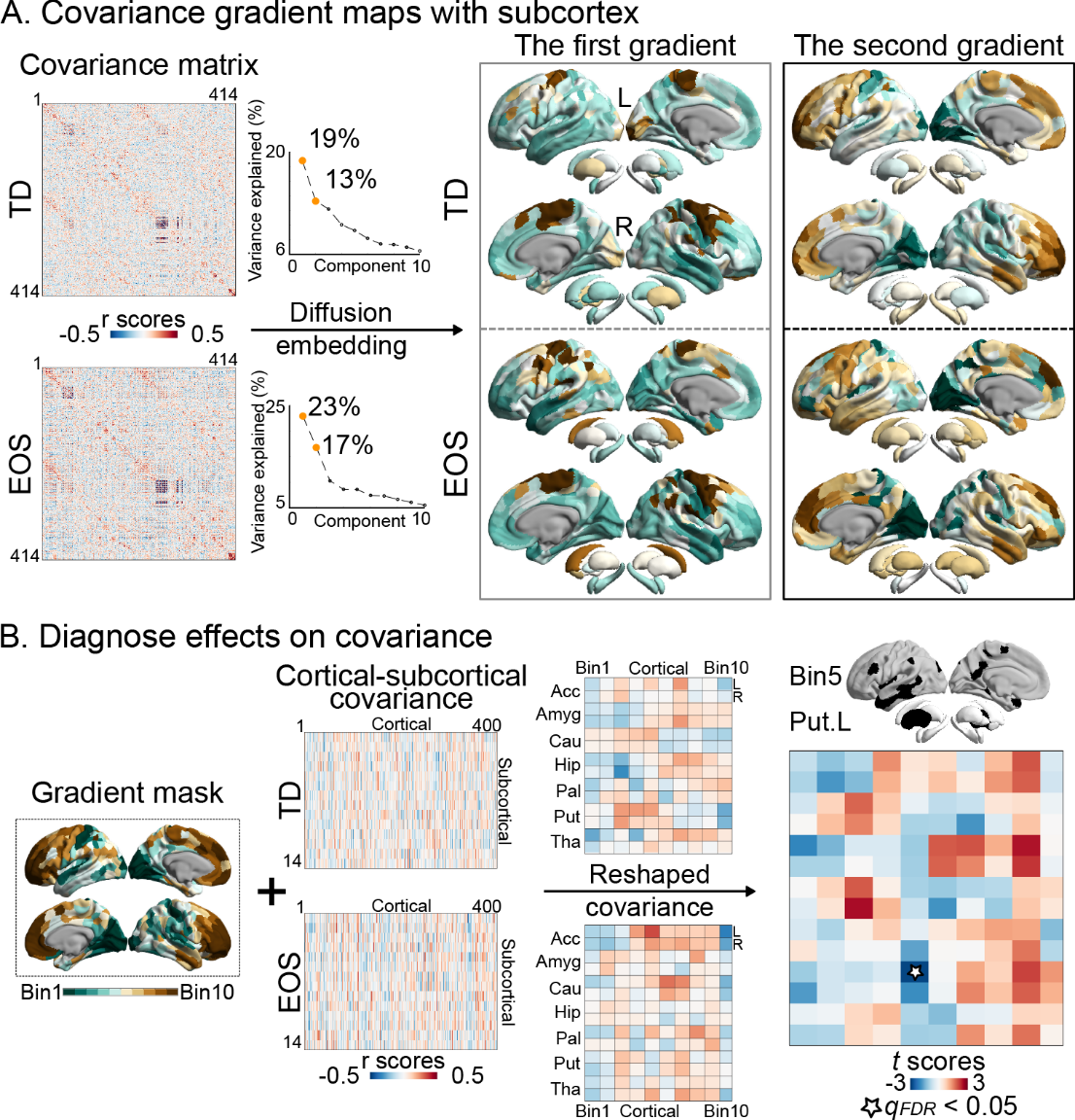

**Figure S5. Subcortical results.** **(A)** Covariance gradients by combining cortex and 14 subcortical nuclei. The first and second gradient maps are similar to the original cortical gradient maps without subcortex. Patients showed visually increased gradient values in anterior and superior nuclei compared to controls. For example, the caudate along the first gradient axis and the putamen along the second gradient axis. **(B)** Disease effects on the reshaped cortical-subcortical covariance matrix. The original normative second gradient mask in TD was used as a mask to reshape the cortical-subcortical covariance matrix. We found that patients had decreased covariance values between the 5th bin and the left putamen (*t* = −2.76, *q_FDR_* = 0.03), i.e. less positive covariance values in EOS compared to TD. Acc, accumbens; Amyg, amygdala; Cau, caudate; Hip, hippocampus; Pal, pallidum; Put, putamen; Tha, thalamus.
